# Yolk platelets impede nuclear expansion in *Xenopus* embryos

**DOI:** 10.1101/2021.01.13.426473

**Authors:** Sora Shimogama, Yasuhiro Iwao, Yuki Hara

## Abstract

During metazoan early embryogenesis, the intracellular properties of proteins and organelles change dynamically through rapid cleavage. In particular, a change in the nucleus size is known to contribute to embryonic development-dependent cell cycle and gene expression regulation. Here, we compared the nuclear sizes of various blastomeres from developing *Xenopus* embryos and analyzed the mechanisms that control the nuclear expansion dynamics by manipulating the amount of intracellular components in a cell-free system. There was slower nuclear expansion during longer interphase durations in blastomeres from vegetal hemispheres than those from animal hemispheres. Furthermore, upon recapitulating interphase events by manipulating the concentration of yolk platelets, which are originally rich in the vegetal blastomeres, in cell-free cytoplasmic extracts, there was slower nuclear expansion and DNA replication as compared to normal yolk-free conditions. Under these conditions, the supplemented yolk platelets accumulated around the nucleus in a microtubule-dependent manner and impeded organization of the endoplasmic reticulum network. Overall, we propose that yolk platelets around the nucleus reduce membrane supply from the endoplasmic reticulum to the nucleus, resulting in slower nuclear expansion in the yolk-rich vegetal blastomeres.

## INTRODUCTION

In the early stages of metazoan embryogenesis, blastomeres divide rapidly to increase the cell number and define cell fates during later development. To generate blastomeres with different cell fates, different cell fate determinants, including proteins, mRNA, and organelles, need to be segregated between the sister blastomeres during cleavage. An asymmetric cell division (cleavage) results in different level of these determinants in different-sized sister cells, as seen in *Caenorhabditis elegans* embryos (Rose & Gönczy, 2014). Additionally, even if the cell divides symmetrically, heterogeneity in the distribution of the cytoplasmic components contributes to a difference in the determinant amount. In metazoan oocytes, some cytoplasmic components and intracellular structures are unevenly distributed in the cytoplasm. In giant oocytes of frog *Xenopus*, proteins including transcription factors and intracellular structures including yolk platelets and pigment granules, are distributed more towards the vegetal hemisphere of the oocytes. In particular, heavy yolk platelets in the cytoplasm and large nucleoli in the nucleus sediment towards the vegetal side by gravitational forces (Neff et al, 1984; Feric & Brangwynne, 2013). Conversely, the yolk-free cytoplasm is located on the side of the animal hemispheres, around the germinal vesicle (GV) (Imoh, 1995). Thus, various determinants such as yolk platelets (Danilchik & Gerhart, 1987), transcription factor VegT (Fukuda et al, 1990), and aPKC (Chalmers et al, 2003) are distributed differently inside the oocytes and segregate to cleaved blastomeres in different amounts. Consequently, during early embryogenesis, a mosaicism of the cytoplasmic determinants is established in the blastomeres, especially between blastomeres that originate from animal and vegetal hemispheres.

During the establishment of embryonic cell fates, there is a drastic change in the morphology of the intracellular structures as well as in the amount of cytoplasmic determinants. The size of the nucleus is one of the important features. During the early stages of metazoan embryogenesis, there is a reduction in the sizes of the nucleus and cell as blastomeres divide without an increase in the cell size. However, the reduction in nuclear size is relatively small, as compared to that in cell size, resulting in an increase in the ratio between cell and nucleus size (N/C ratio) as embryogenesis proceeds (Wesley et al, 2020). At the midblastula stage of *Xenopus laevis*, the increase in N/C ratio is generally stalled and it remains a constant thereafter at later embryonic stages (Jevtić and Levy, 2015). Simultaneously, the timing of reaching this specific N/C ratio is coupled to stages known as midblastula transition (MBT), when the cell cycle changes from synchronous rapid cell division without gap phases to asynchronous cell division (Newport & Kirschner, 1982) and zygotic gene activation (ZGA), when the zygotic genes that are originally repressed during early embryogenesis are activated (Gentsch et al, 2019; Chen et al, 2019a). In addition to these consistencies in the timings, upon experimentally manipulating the nuclear size through genetic perturbations of nuclear envelope components or generation of haploid embryos in *X. laevis* embryos, the timings of MBT and ZGA could be shifted until a certain N/C ratio was achieved (Jevtić and Levy, 2015; Jevtić and Levy, 2017). Furthermore, some cellular functions other than developmental aspects are known to correlate with nuclear size in a wider range of eukaryotes (Edens et al, 2013). Thus, nuclear size is expected to contribute to various intracellular aspects that control embryonic development.

The mechanisms for controlling nuclear size have been well analyzed and can be classified into two concepts: by controlling through cytoplasm (‘external mechanism’) or through the DNA inside the nucleus (‘internal mechanism’). Each concept is based on observations of the constant N/C ratio among cell types in individual organisms and diverse species (Neumann & Nurse, 2007; Jorgensen et al, 2007; Jevtić & Levy, 2015; Price et al., 1973; Hara, 2020) or a positive correlation between nuclear size and DNA content among species and cells with different ploidy (Fankhauser, 1945; Robinson et al, 2018; Gillooly et al, 2015; Hara, 2020). In metazoan embryogenesis, the nuclear size is basically controlled by external mechanisms because there are no significant changes in the DNA amount. Indeed, analyses using in vivo *Xenopus* embryos and a cell-free system of *Xenopus* egg extracts have revealed that the availability of nuclear components including lipid membranes and nuclear lamina components in the cytoplasm determines the plateau size and expansion dynamics of the nucleus in interphase (Hara & Merten, 2015; Mujurree et al, 2020; Levy & Heald, 2010; Edens & Levy, 2014; Edens et al, 2017; Chen et al, 2019b; Brownlee & Heald, 2019). The nuclear lamina components are supplied via nuclear import through nuclear pore complexes (NPCs), whose amount correlates with the cell volume (Levy & Heald, 2010). Lipid membranes are supplied from the cytoplasm to the nuclear membrane, which correlates with the size of the perinuclear endoplasmic reticulum (ER) (Hara & Merten, 2015; Mukherjee et al, 2020). However, these studies have utilized either developmental embryos without separating the blastomeres or homogeneous cell-free cytoplasmic extracts. Therefore, the effects of different cytoplasmic properties in each blastomere on the determination of nuclear size in interphase have not been considered enough. In this study, to evaluate these effects experimentally, we compared the nuclear expansion dynamics among various blastomeres (including vegetal blastomeres with many yolk platelets from *Xenopus* embryos) and evaluated the effects of yolk platelets by manipulating their amount in a cell-free system.

## RESULTS

### The ratio between nucleus and cell volume is not different among animal and vegetal blastomeres from *Xenopus* developing embryos

To observe the nuclei in individual blastomeres from developing embryos, we dissociated the blastomeres from *X. laevis* embryos at various developmental stages. In the process of blastomere dissociation, we manually separated blastomeres into three groups according to superficial features: (1) animal blastomeres with black surface pigments, (2) animal blastomeres without pigments, and (3) vegetal blastomeres. The nucleus was visualized by carrying out immunohistochemistry of NPCs and staining DNA in each blastomere group (Fig. 1A). From the microscopic images, we measured cross-sectional areas of the nucleus and cell, and calculated their volumes. Since the cell cycle was not synchronized spontaneously among the blastomeres and embryos, our measured values included those from various timings of interphase, even when using blastomeres from a single embryo. Corroboratively, when plotting the calculated nuclear volume against the cell volume in a log-log plot, the nuclear volumes revealed variations even in blastomeres with almost the same cell volumes (longitudinal axis in Fig. 1B). Nonetheless, the plot generated using data without blastomere grouping revealed a positive correlation between the calculated nuclear volume and cell volume, as described previously (Jevtić & Levy, 2015) (Fig. 1B). From the plot, we detected a ‘hypoallometric’ relationship, which is indicated as *(nuclear volume) = A × (cell volume) B* where scaling exponent *B* is *< 1*. It should be noted that the calculated scaling exponent from the early developing embryos before MBT was significantly smaller than that from the developed embryo after MBT (Fig. S1A). Upon separating the data of cell and nuclear volumes based on the three blastomere groups, the correlation of nuclear volume with cell volume revealed no substantial differences between the groups (Fig. 1C). Additionally, upon comparing the data of specific animal and vegetal blastomeres with similar cell volumes, the nuclear volumes were not found to be substantially different (Fig. 1D). These results suggest that the N/C ratio does not vary, regardless of the blastomere group. It should be noted that these observed features of nuclear size scaling were also observed in dissociated blastomeres from embryos of sister species *Xenopus tropicalis* (Fig. S1B–D), suggesting that these features are conserved within at least the *Xenopus* species.

**Figure 1.**
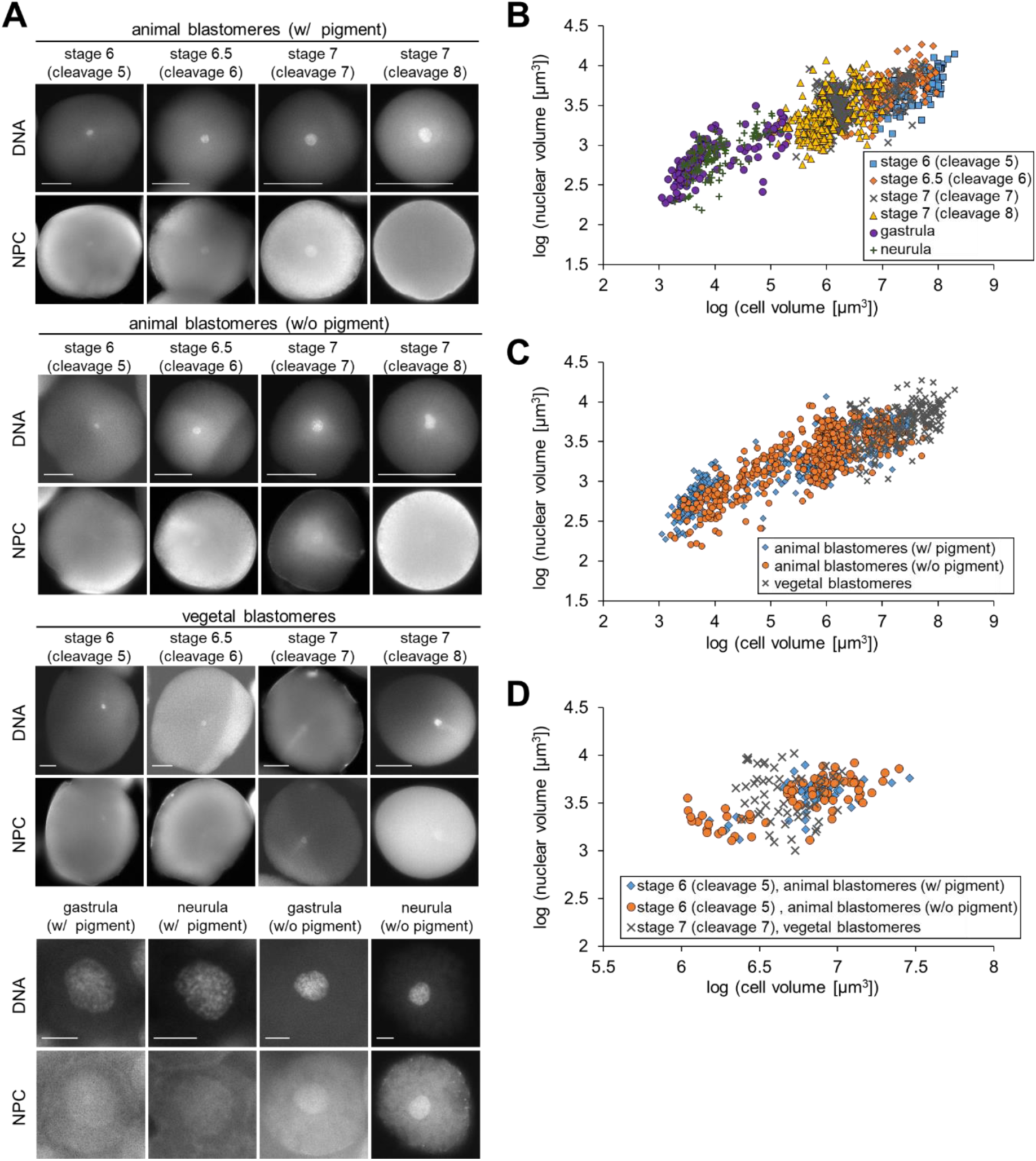
Constant N/C ratio among animal and vegetal blastomeres from developing *X. laevis* embryos. (**A**) DNA (top; stained using TO-PRO-3) and NPCs (bottom: immunostaining) were visualized in dissociated blastomeres from *X. laevis* developing embryos. The blastomeres from embryos at various developmental stages were classified into three different groups; blastomeres with black pigments from the animal hemisphere [animal blastomeres (w/ pigments)], blastomeres without black pigments from the animal hemisphere [animal blastomeres (w/o pigments)], and blastomeres from vegetal hemispheres (vegetal blastomeres). Scale bar: 100 µm. The animal blastomeres from gastrula and neurula embryos were classified into those with or without pigments. Scale bar: 10 µm. (**B**) Calculated nuclear volume was plotted against the calculated cell volume of blastomeres from different developmental stages of embryos. (**C**) Data of the nuclear and cell volumes were classified into the different blastomere groups (different colors). The values were identical to those in panel B. (**D**) Only data of the nuclear and cell volumes in animal blastomeres from stage 6 (cleavage 5) embryos and vegetal blastomeres from stage 7 (cleavage 7) embryos were represented.

### Nucleus expands slowly in vegetal blastomeres

During early stages of embryogenesis in *Xenopus*, the nucleus expands continually over the interphase and cannot reach a plateau volume within the limited duration of the interphase (Levy & Heald, 2010; Heijo et al, 2020). To confirm this feature of continuous nuclear expansion among the three different blastomere groups, we analyzed the nuclear expansion dynamics under extended interphase by adding cycloheximide to block the spontaneous transition to mitosis (Fig. 2A). In all the blastomere groups, the calculated nuclear volume was larger than that in the normal condition at each developmental stage (Fig. 2A and 2B), suggesting that the nucleus expands continually in the extended interphase, regardless of the blastomere groups. It should be noted that the extended nuclear expansion was only observed until 3 h of incubation after addition of cycloheximide, which might have been caused by the induction of apoptotic phenomenon through long-term exposure of the embryos to cycloheximide (Gibeaux et al, 2018). Next, to compare the nuclear expansion speed among the blastomere groups, we measured the duration of the cell cycle and considered it as the speed of nuclear expansion because it is technically difficult to directly measure the nuclear expansion speed through time-lapse observation in opaque blastomeres. By incubating the dissociated blastomeres under a stereoscopic microscope, we measured the duration from the completion of one cytokinesis to the next (Fig. 2C). As a result, when comparing the length of cell cycle duration in vegetal blastomeres to that in animal blastomeres exhibiting almost the same cell volume, the length was remarkably longer in vegetal blastomeres than in animal blastomeres (see the difference at the longitudinal axis in Fig. 2C). Conversely, the duration from initiation to completion of cleavage furrow formation was not significantly different between animal and vegetal blastomeres (Fig. S2). These results suggest that the cell cycle duration (mostly in interphase, rather than the whole cell cycle duration) is longer in vegetal blastomeres than in animal blastomeres. It should be noted that the detected cell cycle duration in each animal and vegetal blastomere group correlated negatively with cell volume (Fig. 2C), which is consistent with previous observations involving only animal blastomeres (Iwao et al, 2005). The detected longer cell cycle in vegetal blastomeres and the constant N/C ratio between animal and vegetal blastomeres indicate that the nucleus expands at a much slower rate in vegetal blastomeres and this expansion continues until it reaches the same N/C ratio as the animal blastomeres.

**Figure 2.**
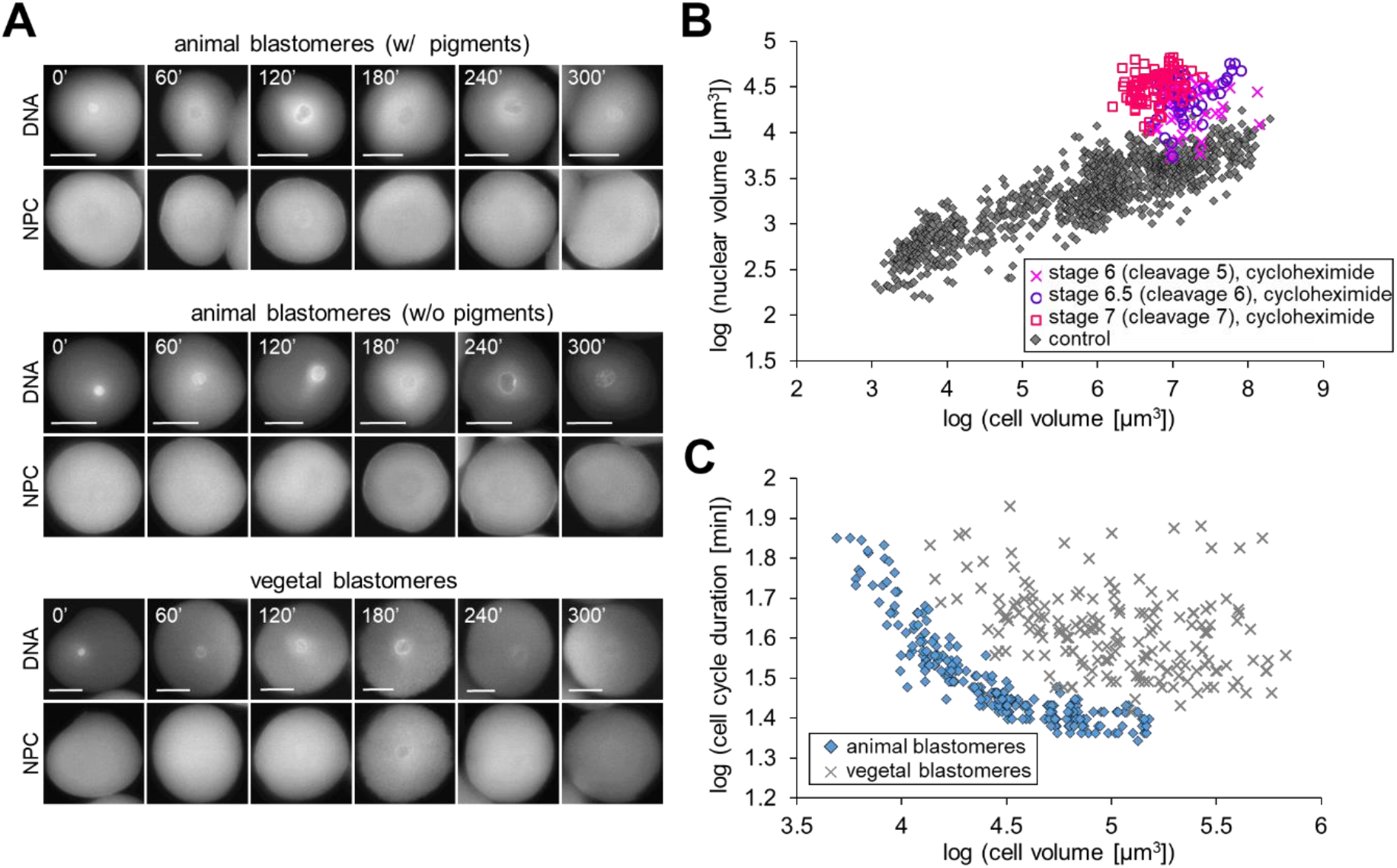
Nuclear expansion in animal and vegetal blastomeres from *X. laevis* embryos. (**A**) DNA (top: stained using TO-PRO-3) and NPCs (bottom: immunostaining) were visualized in dissociated blastomeres of three different groups from *X. laevis* embryos treated with cycloheximide. Time represents that after incubation of the dissociated blastomeres with cycloheximide. Scale bar: 100 µm. (**B**) Calculated nuclear volume was plotted against the calculated cell volume of the blastomeres treated with cycloheximide for 180-240 min (color symbols). The data (grey) in the control condition without cycloheximide were identical to those in Fig. 1B. (**C**) Measured duration of a single cell cycle, from the completion of one cytokinesis to the next, was plotted against the calculated mean cell volume during the cell cycle. The data were acquired from each dissociated animal (blue) or vegetal (grey) blastomere.

### Existence of yolk platelets decreases the speed of nuclear expansion and interphase progression

In *X. laevis* oocytes, various cytoplasmic components, including proteins and organelles, are distributed in a biased manner along the animal-vegetal axis. Yolk platelets, which are remarkably biased components in oocytes, are segregated more towards vegetal blastomeres through embryonic cleavages. To assess whether a difference in the concentration of yolk platelets among the blastomere groups would contribute to the nuclear expansion dynamics, we first experimentally manipulated their concentration in animal blastomeres *in vivo*. By carrying out mild centrifugation just before the 3^rd^ cleavage along the equatorial plane in the embryos, the yolk platelets were sedimented to the vegetal side, while translucent blastomeres with less yolk platelets were generated after a few cleavages on the animal side of the embryos (Fig. 3A). Observation of the nucleus in the blastomeres revealed that the measured nuclear volumes in translucent animal blastomeres were not substantially different from those in normal animal blastomeres with almost the same cell volume (Fig. 3B). This result suggests that yolk platelets and other components sedimented using centrifugation do not contribute much to determine the nuclear expansion dynamics in the animal blastomeres. Nonetheless, since animal blastomeres are expected to have a relatively lower concentration of yolk platelets, as compared to vegetal blastomeres, their concentration in animal blastomeres is possibly less effective in changing the nuclear expansion dynamics. Next, we recapitulated the cytoplasmic condition with a much higher concentration of yolk platelets using a cell-free reconstruction system, since it was technically difficult to handle vegetal blastomeres containing many sedimented yolk platelets during the preparation of translucent blastomeres. We isolated yolk platelets from *X. laevis* eggs (Fig. 3C) and supplemented them at three different concentrations (final concentrations: high, 1-2×10^5^/µl; medium, 2-4×10^4^/µl; low, 4-8×10^3^/µl) into yolk-less cytoplasmic extracts from *X. laevis* eggs. First, we supplemented the isolated yolk platelets and *X. laevis* sperm chromatin into the interphase extracts for reconstructing the nuclei (Fig. 3D). In the presence of yolk platelets, spherical nuclei enclosed by membranes were successfully reconstructed and yolk platelets were detected around the nuclei [yolk platelets strongly labeled using DiOC_6_(3) in Fig. 3D]. Using these samples, we measured the cross-sectional area of the reconstructed nuclei at each incubation time point and analyzed the nuclear expansion dynamics (Fig. 3E). The observed nuclear expansion was significantly slower in the presence of yolk platelets at high and medium concentrations than in the buffer control (Fig. 3E). Interestingly, the resulting nuclear expansion speed decreased in a dose-dependent manner in the supplemented yolk platelets (Fig. 3E). However, the nuclear expansion in the presence of yolk platelets supplemented at low concentrations was not substantially different from that in the control condition. This is consistent with our observation that there is less difference in the nuclear size between yolk-less translucent blastomeres and normal animal blastomeres originally containing small amounts of yolk platelets (Fig. 3B). It should be noted that the observed nuclear cross-sectional area varied among individual preparations of cytoplasmic extracts. Therefore, we set criteria that would allow us to use the measured data for analyses in this study (see Materials and Methods), and also confirmed the nuclear expansion dynamics using normalized values of the nuclear cross-sectional area for each extract preparation (Fig. S3A). Additionally, to analyze the effect of yolk platelets on DNA replication, we visualized the incorporation of fluorescently-labeled nucleotides into the chromatin and measured the time taken to complete DNA replication. As a result, the completion timing was delayed only in the condition supplemented with yolk platelets at the highest concentration (Fig. 3F). To further evaluate whether the existence of yolk platelets can influence cell cycle events, we measured the duration of the cell cycle by utilizing a cycling extract (Fig. 3G). Generally, this extract supplemented with sperm chromatin can initiate interphase and progress spontaneously through the M−S−M cell cycle several times. As a result, the duration of 1^st^ interphase from initiation of the incubation to the nuclear envelope break down (NEBD) was significantly longer upon supplementation of yolk platelets at a high concentration into the cycling extract than in the buffer control (Fig. 3G). In contrast, the duration of mitosis, from NEBD to the assembly of membrane-enclosed spherical chromatin, was not different among the groups (Fig. 3G), which is consistent with the observation that there was no significant difference in the duration of cleavage furrow formation between *in vivo* animal and vegetal blastomeres (Fig. S2). Collectively, the presence of yolk platelets at high concentrations in the cytoplasm resulted in slowed interphase progression, including nuclear expansion and DNA replication.

**Figure 3.**
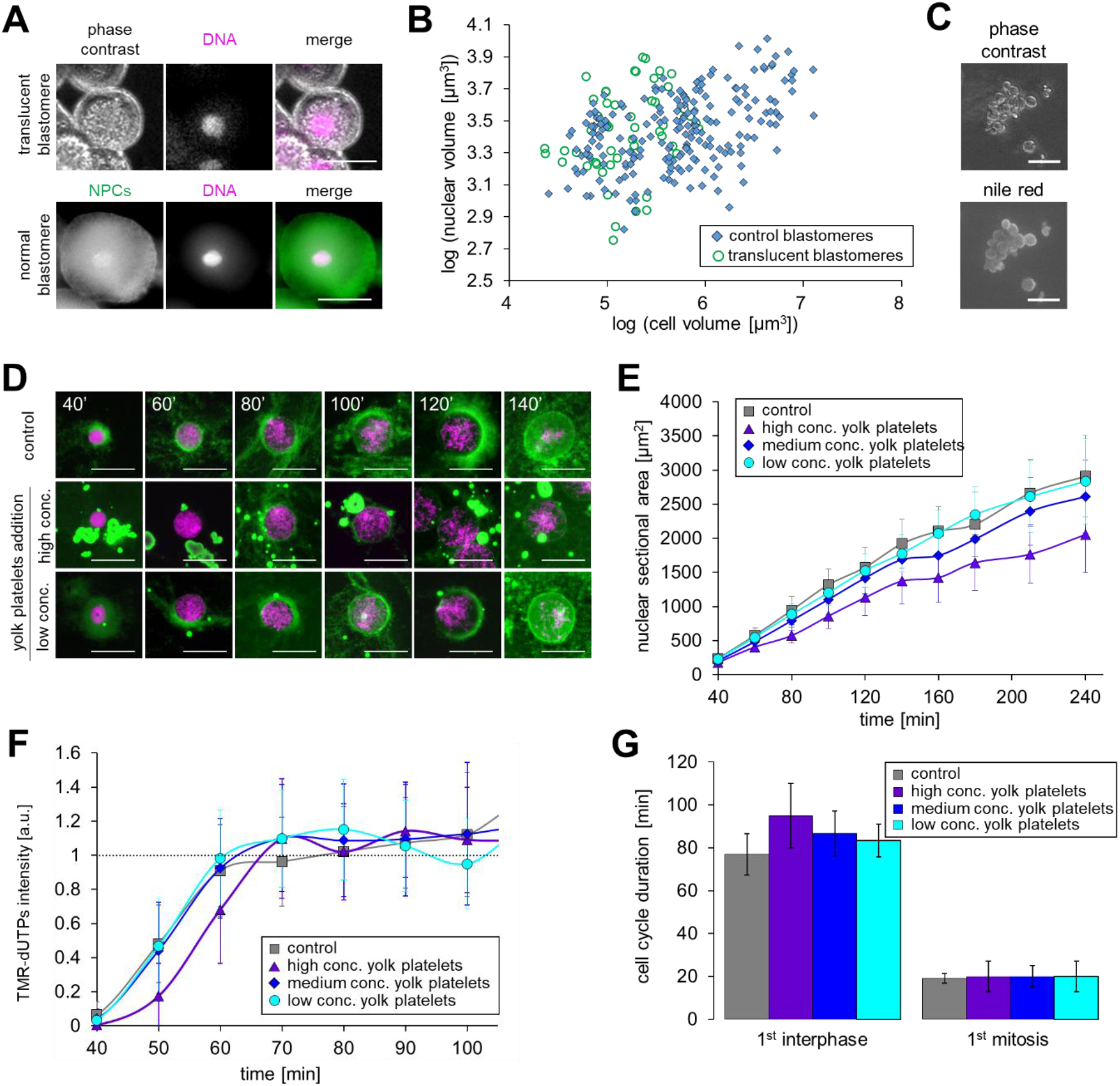
Supplementation with yolk platelets in cell-free extracts decreases the speed of interphase progression. (**A**) Representative images of translucent (top) and normal (bottom) blastomeres from *X. laevis* embryos. DNA (magenta) was visualized using Hoechest33342 and merged with phase contrast image (top) or an image of NPCs (green), visualized using immunohistochemistry (bottom). Scale bar: 50 µm. (**B**) Calculated nuclear volume was plotted against the calculated cell volume of translucent (green circle) and control dissociated (blue diamond) blastomeres. The data of each blastomere were obtained from embryos from the same mother frog. (**C**) Representative images of isolated yolk platelets - phase contrast (top) and stained using Nile Red (bottom). Scale bar: 30 µm. (**D**) Representative images of reconstructed nuclei in *X. laevis* egg extracts upon supplementation with isolated yolk platelets at different concentrations. DNA (magenta; stained using Hoechst33342) and membrane [green; stained by DiOC_6_(3)] were visualized in the fixed sample at the indicated time of incubation. Scale bar: 50 µm. (**E**) Dynamics of the measured mean cross-sectional area in the nuclei reconstructed with different concentrations of yolk platelets (control: n = 9; high (1-2×10^5^/µl): n = 6; medium (2-4×10^4^/µl): n = 6; low (4-8×10^3^/µl): n = 5). (**F**) Dynamics of the intensities of the incorporated TMR-dUTPs within the nucleus. The intensity was calculated by multiplying the measured TMR-dUTPs intensity (/µm^2^) with the measured nuclear cross-sectional area. Each calculated value was divided by the mean value of the individual extract preparation after 120 min of incubation in the control condition. control: n = 7; high conc.: n = 4; medium conc.: n = 4; low conc.: n = 3. Averages of values from each extract preparation are connected using a line in each dataset. (**G**) Measured duration of each cell cycle upon supplementation with the isolated yolk platelets at different concentrations in the cycling extract. The extracts, upon addition of sperm chromatin, initially expose interphase (1^st^ interphase), then transit spontaneously to mitosis (1^st^ mitosis), and interphase of the next cell cycle. Averages (±SD) from each extract preparation were shown in each dataset. control: n = 8; high conc.: n = 3; medium conc.: n = 3; low conc.: n = 3.

### Yolk platelets physically impede the nuclear expansion process

The observed decrease in the nuclear expansion speed upon supplementation with yolk platelets allows us to consider two possibilities: either the total volume or the total number of yolk platelets might contribute to the nuclear expansion. To evaluate this, we separated roughly small-sized and large-sized yolk platelets populations from the bulk isolated solution by further fractionation (Fig. 4A and 4B) and supplemented with either of the fractions at the same concentration (1-2×10^5^/µl) into the interphase extracts (Fig. 4C). Supplementation with the large-sized faction into the extract resulted in slower expansion of the reconstructed nuclei, as compared to supplementation with the bulk fraction of yolk platelets (Fig. 4D). On the contrary, upon supplementation with the small-sized fraction, the nuclear expansion was more rapid than that observed in case of the bulk fraction, but slower than that in the control condition (Fig. 4D). It should be noted that the dependency of nuclear expansion speed on the size of yolk platelets was also observed using normalized values of the nuclear cross-sectional area for each extract preparation (Fig. S3B). Furthermore, to comprehensively understand all the results obtained upon supplementation with yolk platelets of different sizes and concentrations, we calculated the total volume and volume occupancy of the yolk platelets that were supplemented into the extract. When the calculated volume occupancies of yolk platelets were plotted against the calculated speed of nuclear expansion, a negative correlation was observed (Fig. 4F). These data indicate that a reduction in nuclear expansion speed is dependent on the total volume of yolk platelets in the cytoplasm rather than on the number of supplemented yolk platelets.

**Figure 4.**
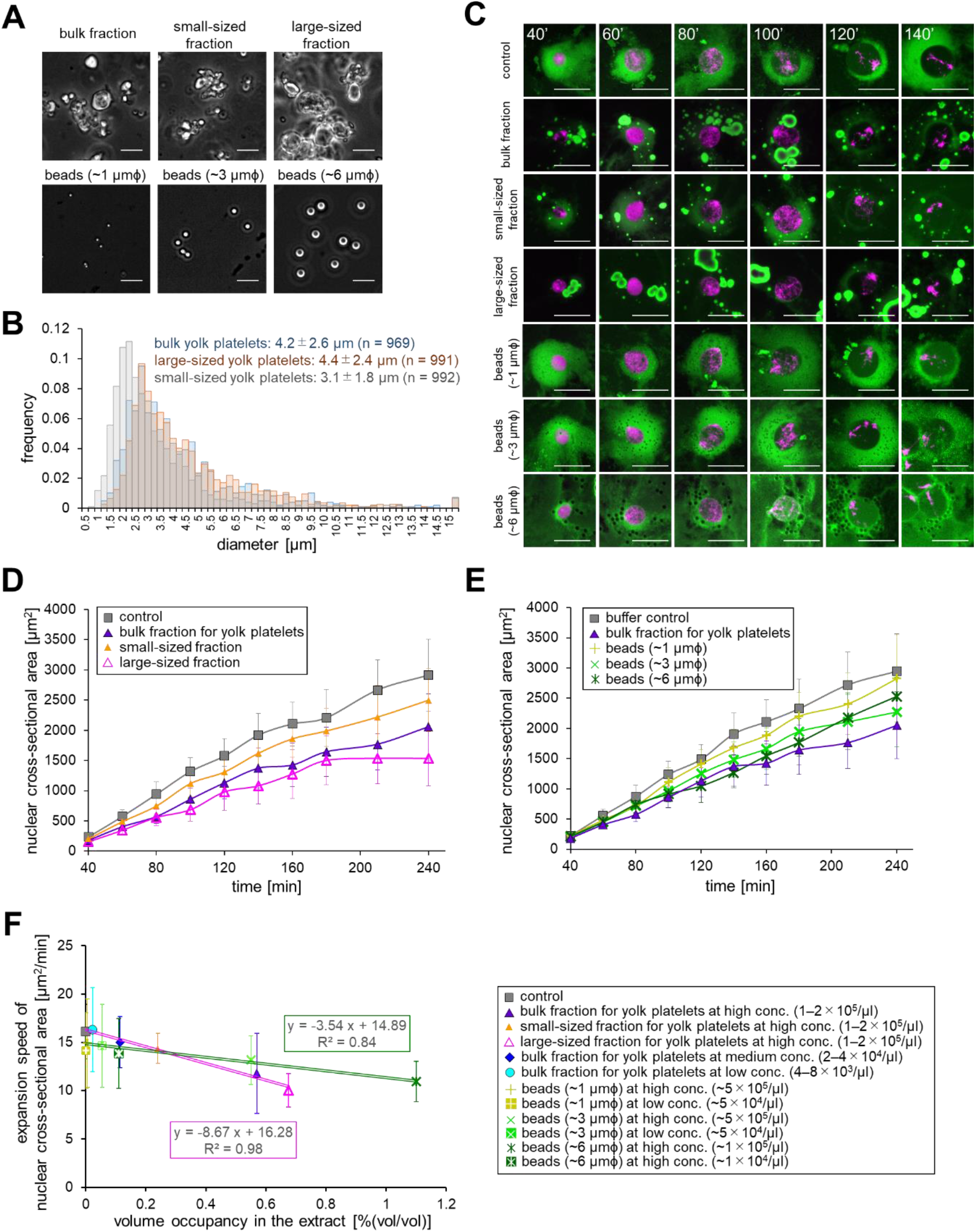
Nuclear expansion speed depends on the total volume of yolk platelets in the cytoplasm. (**A**) Representative images of isolated yolk platelets in bulk fraction, small-sized fraction, large-sized fraction, and images of 1 µm diameter magnetic beads, 3 µm diameter magnetic beads, and 6 µm diameter polystyrene beads. Scale bar: 10 µm. (**B**) Diameter distributions of each fraction of yolk platelets. (**C**) Representative images of reconstructed nuclei in *X. laevis* egg extracts upon supplementation with either fraction for yolk platelets or artificial beads. DNA (magenta; stained using Hoechst33342) and membrane [green; stained by DiOC_6_(3)] were visualized in the fixed sample at the indicated time of incubation. Scale bar: 50 µm. (**D**) Dynamics of the measured mean cross-sectional area in the reconstructed nuclei by supplementation with different-sized yolk platelets at a concentration of 1-2×10^5^/µl (control: n = 9; bulk fraction for isolated yolk fraction: n = 5; small-sized fraction: n = 3; large-sized fraction: n = 3). (**E**) Dynamics of the measured mean cross-sectional area in the reconstructed nuclei upon supplementation with each artificial beads [1 µm diameter beads (∼5×10^5^/µl): n = 4; 3 µm diameter beads (∼5×10^5^/µl): n = 6; 6 µm diameter beads (∼1×10^5^/µl)] or controls [buffer control: n = 9; bulk fraction for yolk platelets (1-2×10^5^/µl): n = 6; which are identical to those in panel D]. Averages of values from each extract preparation are connected using a line in each dataset. (**F**) Calculated expansion speed of the nuclear cross-sectional area was plotted against the volume occupancy of yolk platelets or artificial beads in the cytoplasmic extract. Average values from each extract preparation are plotted. Each data set, upon supplementation with yolk platelets (pink) or artificial beads (green) is fitted using linear regression. Error bar: SD.

Next, to assess if components other than yolk platelets can reduce the nuclear expansion speed, we utilized artificial alternatives instead of isolated yolk platelets. We reconstructed the nuclei by supplementation with various concentrations of magnetic beads with diameters of 1 µm or 3 µm, or polystelene beads with diameters of 6 µm (Fig. 4A). Interestingly, in the presence of either kind of artificial beads, the nuclei were successfully reconstructed and many beads were detected around the nucleus (black circles in the fluorescent images in Fig. 4C). Under these conditions of supplementation with artificial beads, especially at much higher concentrations, there was a decrease in the nuclear expansion speed, as compared to that in the control condition (Fig. 4E). Furthermore, the decrease in nuclear expansion speed could be detected by supplemented with different concentrations in cases of the 6 µm diameter beads (at a concentration of ∼1×10^5^/µl) and 3 µm or 1 µm diameter beads (at a concentration of ∼5 × 10^5^ /µl) (Fig. 4E), suggesting that the total volume of beads supplemented into the extracts contributes to the nuclear expansion control. Corroboratively, the calculated volume occupancies of the artificial beads correlated negatively with the calculated nuclear expansion speed under conditions of supplementation with different-sized beads at different concentrations (Fig. 4F). It should be noted that in case of artificial beads, the same dependency was also observed when using normalized values of the nuclear cross-sectional area for each extract preparation as well (Fig. S3C). Although the dependency of nuclear expansion dynamics on volume occupancy was conserved between yolk platelets and artificial beads, the slope of the regression line of the nuclear expansion speed with the volume occupancy of the components was steeper in case of yolk platelets than in case of artificial beads (Fig. 4F). Furthermore, when artificial beads were supplemented at a concentration that resulted in effective slow nuclear expansion with yolk platelets, there was less delay in the completion timing of DNA replication (Fig. S3D). These results suggest that yolk platelets are more effective than artificial beads in reducing the nuclear expansion speed and DNA replication. Collectively, the existence of micrometer-sized structures in the cytoplasm could reduce the nuclear expansion speed in a volume-occupancy-dependent manner.

### Yolk platelets impede the organization of the endoplasmic reticulum

Considering that the supplemented yolk platelets and artificial beads were detected around the nucleus (Fig. 3D and 4C), we hypothesized that these micrometer-sized structures can influence the functional space surrounding the nucleus to control nuclear expansion. During the nuclear formation and expansion phase, lipid membranes accumulate around the chromatin and are supplied to the nuclear membrane through the ER around the nucleus. The organization of functional ER networks is controlled by microtubules (MTs) radiating from the centrosome, in association with motor proteins (Reinsch and Karsenti, 1997; Niclas et al, 1996). To determine how yolk platelets influence the organization of the ER networks on MTs, we first observed the dynamics of yolk platelets during nuclear formation and expansion in the extracts. Through time-lapse imaging of the extracts, the supplemented yolk platelets, which were initially distributed evenly in the cytoplasmic extracts, moved toward the chromatin (Fig. 5A). After approximately 40 min of incubation in the extract, many yolk platelets accumulated around the small assembled nucleus, which is consistent with our observation of yolk platelets located around the nuclei on fixed samples (Figs. 3D and 4C). Accordingly, many yolk platelets and artificial beads were detected in the MT-occupied space in the fixed samples sedimented on a glass slide (Fig. 5B). In the presence of nocodazole, an inhibitor for MT polymerization, in the extract, the yolk platelets rarely moved toward the chromatin (Fig. 5A), suggesting that yolk platelets move toward the minus-end of MTs. However, the size of MT-occupied space itself showed less difference among samples that were/were not supplemented with yolk platelets or artificial beads (Fig. 5C). The accumulation of yolk platelets or artificial beads was possibly achieved through non-specific attachment of floating yolk platelets in the extract to the glass slide during the sedimentation step of the observation slide preparation. However, yolk platelets and artificial beads were rarely detected in the region of the glass slide where MTs were not sedimented (Fig. 5B), suggesting that the detected yolk platelets are located within the MT-occupied space before preparation of the observation slide. Furthermore, we used the images obtained from samples with different concentrations of yolk platelets and magnetic beads to calculate the area occupancy of yolk platelets or beads within the area of the MT-occupied space (Fig. 5D). The estimated area occupancy increased with an increase in the concentration of the supplemented yolk platelets (Fig. 5D) and correlated negatively with the observed nuclear expansion speed (Fig. 3E). Additionally, upon comparing the conditions of supplementation with yolk platelets and magnetic beads, which showed almost the same volume occupancies within the cytoplasmic extracts (i.e., bulk fraction of yolk platelets at high concentration and 3 µm diameter beads at high concentration), the detected area occupancies within the MT-occupied space were significantly higher in case of yolk platelets (Fig. 5D). This suggests that the artificial beads interact with MTs in a weaker fashion, as compared to the yolk platelets, resulting in a gentler observed slope of the correlation between nuclear expansion speed and volume occupancy within the extracts (Fig. 4F). Finally, when we visualized ER networks in the MT-occupied space using the specific fluorescent dye DiIC_16_(3) during the nuclear expansion phase, the ER meshwork was less organized in the vicinity of yolk platelets, which appeared to generate holes on the ER meshwork (arrowheads in Fig. 5E). This result suggests that the yolk platelets in the MT-occupied space physically impede the organization of the ER networks. This disorganization of the ER networks in the MT-occupied space could reduce the ability of the lipid membrane supply from the ER to the nucleus in an occupancy-dependent manner, resulting in a decrease in nuclear expansion speed in the presence of yolk platelets or artificial beads.

**Figure 5.**
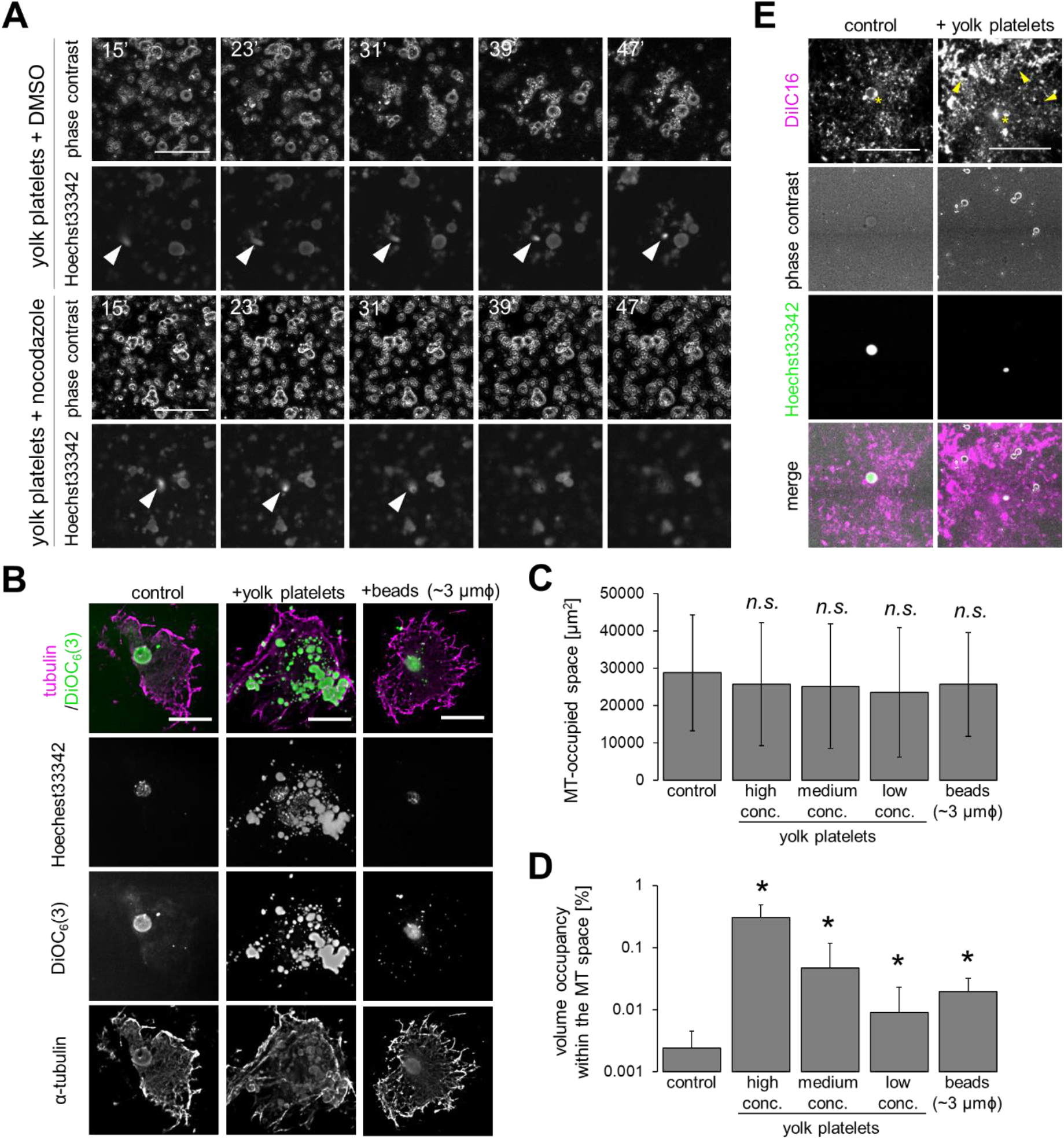
Yolk platelets impede organization of the ER within the MT-occupied space. (**A**) Snapshots of representative time-lapse images of the reconstructed nuclei in the cytoplasmic extract upon supplementation with yolk platelets at a concentration of 1-2×10^5^/µl in the presence of DMSO (for buffer control: top) or nocodazole (bottom). DNA was visualized using Hoechest33342. Bright field images were taken using phase contrast. Arrowheads represent chromatin in the extracts. Scale bar: 50 µm. (**B**) α-tubulin (magenta; immunostaining), membranes [green; stained by DiOC_6_(3)], and DNA (blue; stained using Hoechst33342) were visualized in the sedimented samples on a glass slide. The nuclei were reconstructed by supplementation with buffer (control), yolk platelets at a concentration of 1-2×10^5^/µl (high concentration), or 3 µm diameter magnetic beads at a concentration of ∼5×10^5^/µl for 120 min of incubation. Scale bar: 100 µm. (**C**) Measured mean area of MT-occupied space in each sample upon supplementation with buffer (n = 35), yolk platelets at high concentration (n = 32), medium concentration (2-4×10^4^/µl: n = 36), low concentration (4-8×10^3^/µl: n = 31), or 3 µm diameter magnetic beads (n = 41). (**D**) Calculated mean area occupancy of yolk platelets or magnetic beads within the MT-occupied space. Error bars, SD. Asterisks, *P* values obtained using Wilcoxon test, compared to the control condition (*P* < 0.05). *n*.*s*., no significant difference.

## DISCUSSION

*Xenopus* eggs contain a large amount of yolk platelets with a size range of 1-14 µm, which make up 50% of the egg volume (Jorgensen, 2008; Fig. 4B). The yolk platelets are concentrated in the vegetal hemispheres of the eggs and segregate to greater degree into the vegetal blastomeres through embryonic cleavages, which is thought to contribute to cell fate decisions during later development (Danilchik & Gerhart, 1987; Imoh, 1995). In terms of intracellular functions, the biasedly located yolk platelets are known to impede the formation of cleavage furrows during embryonic cleavages (Neff et al, 1984; Newport & Kirschner, 1982). In the present study, we examined the intracellular functions of yolk platelets from a new aspect of nuclear size control through in vivo observations of dissociated blastomeres and experimental manipulation of the concentration of yolk platelets in the cytoplasm. Imaging of dissociated blastomeres revealed that the nucleus in vegetal blastomeres expands to reach almost the same N/C ratio as seen in animal blastomeres, but at a much slower speed (Figs. 1–2). Furthermore, manipulation of yolk platelet concentration using a cell-free reconstruction system revealed that yolk platelets accumulated around the nucleus through MT-based mechanisms (Fig. 5) and impeded interphase progression, including control of cell cycle duration, DNA replication, and nuclear expansion (Fig. 3). Interestingly, in vegetal blastomeres, the slow nuclear expansion is in sync with the slow interphase progression, including intranuclear events of DNA replication, to maintain the same N/C ratio as animal blastomeres with rapid nuclear expansion in the short interphase. The existence of yolk platelets is possibly a key factor in controlling multiple nuclear phenotypes, including nuclear expansion as well as cell cycle progression of the interphase. Nuclear size regulation is known to be involved in the control of ZGA and morphological MBT events at the mid-blastrula stage of *X. laevis* embryos. When the nuclear expansion speed decreases upon experimental manipulation and the resulting N/C ratio remain low, the timings of ZGA and MBT shift at later developmental stages (Jevtić & Levy, 2015; Jevtić & Levy, 2017). This contribution of small nuclear size to embryonic development appears to be applicable to relatively slow differentiation of vegetal blastomeres. A recent study revealed that the timing of ZGA in vegetal blastomeres is delayed into the later developmental stages compared with that in animal blastomeres (Chen et al, 2019a). Therefore, maintaining the low N/C ratio by slow nuclear expansion might contribute in maintaining undifferentiated state, along with repression of zygotic genes in vegetal blastomeres. Although these putative biological significances of yolk accumulation and nuclear expansion control are required for further evaluation, yolk platelets have indispensable roles in controlling intracellular functions including, nuclear size and cell cycle, other than nutrient supply for embryogenesis.

In developing *Xenopus* embryos, the ability to supply nuclear constituents, including lamina and lipid membranes from cytoplasm to nucleus, generally controls the nuclear size and its expansion speed (Levy & Heald, 2010; Hara & Merten, 2015; Brawnlee & Heald, 2019; Mujeehker et al, 2020). Our cell-free experiments combined with the manipulation of yolk platelet concentration clearly indicated that yolk platelets move toward the nucleus along the MTs, and the accumulated yolk platelets around the nucleus impede nuclear expansion through disorganization of ER networks within the MT-occupied space (Figs. 3–5). The experiments carried out using artificial beads instead of the yolk platelets revealed that yolk platelets are physically able to impede the nuclear expansion. In our experiments, we supplemented yolk platelets into the cytoplasmic extracts at a highest concentration of 5×10^6^/µl, which corresponds to ∼1/10 of the estimated concentration of yolk platelets in whole eggs. Due to the technical limitation for handling the isolated yolk platelets with extremely high viscosity, it was not possible to recapitulate the in vivo concentration of yolk platelets in the cell-free extracts. Nevertheless, the relatively high volume-occupancy [> 0.5 %(v/v)] of yolk platelets was able to impede nuclear expansion in our cell-free condition. With respect to the intracellular distribution and movement of yolk platelets, a previous study using an electron microscope provided evidence that yolk platelets appear to accumulate around the nucleus, rather than being evenly distributed in the blastomeres of *X. laevis* developing embryos (Imoh, 1995). Furthermore, MT-dependent movement is consistent with previous observations of the minus-end directed movement of yolk granules in *C. elegans* embryos (Kimura & Kimura, 2010), suggesting that the intracellular accumulation and MTs-dependent movement of yolk platelets might be conserved among metazoan species. In contrast, in *Xenopus* oocytes with a giant GV, the yolk platelets are located less around the GV, while a yolk-less cytoplasm surrounds the GV (Imoh, 1995). This specific distribution of yolk platelets in oocytes might possibly reduce the physical impedance from yolk platelets, which maximizes the nuclear expansion speed for generating a giant GV. Apart from our observed physical impedance of yolk platelets, there are other possibilities; such as, yolk platelets could serve as interaction platforms to interact with cell cycle regulation factors and nuclear size determinants, to decrease their functional amount in the cytoplasm. Indeed, the amount of importin alpha, which regulates nuclear expansion, is modulated through binding to lipids of the plasma membrane, resulting in reduction in the functional amount of importin alpha in the cytoplasm (Brawnlee & Heald, 2019), suggesting that the supplemented yolk platelets might reduce the functional amounts of importin alpha and other determinants of nuclear size and cell cycle in the cytoplasm by binding to them. Another possibility is that the amount of some known accelerators for DNA replication, such as DNA polymerase and cdk2 kinase, which are exclusively distributed in animal hemispheres in eggs (Nagano et al 1982; Iwao et al, 1993) and expected to be segregated more in animal blastomeres after embryonic cleavages, might cause a shorter interphase duration and rapid DNA replication in animal blastomeres. Upon exclusion of the functional cytoplasmic volume by the volume of yolk platelets, there would be a greater decrease in the net amount of these accelerators in the vegetal blastomeres. Thus, our findings open a new avenue for considering the effects of yolk platelets in the blastomeres on intracellular functions other than nutrient supply. It is worth investigating the other intracellular effects of yolk platelets in various aspects, such as other intracellular functions, other organelles, and other organisms, to reveal differential intracellular distribution of yolk platelets and comprehensively understand the mechanisms that control nuclear expansion and cell cycle control in whole in vivo embryos during early embryogenesis.

## Material and methods

### Dissociation of blastomeres from *Xenopus* embryos

Fertilized eggs of *Xenopus laevis* were obtained by insemination *in vitro* of fleshly laid *X. laevis* eggs with crushed *X. laevis* testes. Thirty minutes after insemination, embryos were dejellyed with 2% cysteine (pH 7.8 (KOH)) and washed by 0.2× MMR (1× MMR: 0.25 mM 4-(2-hydroxyethyl)-1-piperazineethanesulfonic acid [HEPES]-KOH, 0.1 mM Ethylenediamine-N,N,N’,N’-tetraacetic acid [EDTA], 5 mM NaCl, 2 mM KCl, 1 mM MgCl_2_, 2 mM CaCl_2_, pH 7.8) several times. For *Xenopus tropicalis* embryos, fertilized eggs were obtained by insemination *in vitro* or by mating and were dejellyed with 3% cysteine (pH 7.8 (KOH)). After incubation of *X. laevis* or *X. tropicalis* embryos at stage 4 (cleavage 3), the embryos were transferred to a Ca-Mg-free medium (CMFM: 88 mM NaCl, 1 mM KCl, 2.4 mM NaHCO_3_, 7.5 mM tris(hydroxymethyl)aminomethane [Tris]-HCl, pH 8.0) for dissociating blastomeres. Some embryos were incubated continually to 0.2× MMR to confirm the developmental stages of the intact embryos. Developmental stages of *X. laevis* and *X. tropicalis* embryos were determined according to Nieuwkoop and Faber’s table (Nieuwkoop & Faber, 1994). When appropriate developmental stages were determined in the intact embryos, fertilization membranes of the embryos with dissociated blastomeres were manually removed using fine forceps. Using a mouth pipette under a dissecting microscope, the dissociated blastomeres were separated into three groups: blastomeres from animal hemispheres (‘animal blastomeres’) with black pigments on the blastomere surface, those without black pigments, and those from vegetal hemispheres (‘vegetal blastomeres’). To arrest the cell cycle of blastomeres in the interphase, embryos with dissociated blastomeres enclosed in the fertilization membrane were incubated with 100 µg/ml cycloheximide (Sigma, C7698) in CMFM. The interphase was arrested in the interphase after going through one cell cycle by supplementing with cycloheximide. After 1 h of incubation with cycloheximide, the fertilization membrane was removed. The dissociated and arrested blastomeres were continuously incubated and subjected to fixation and immunostaining using the following procedures at each hour of incubation post cycloheximide supplementation.

The translucent blastomeres with fewer yolks and pigment granules were prepared as previously described (Iwao et al, 2005; Heijo et al, 2020) with slight modifications. Briefly, the dejellyed *X. laevis* embryos at stage 3 (cleavage 2) just before the cleavage furrow formation at stage 4 (cleavage 3) were floated on 32% Ficoll (wt/vol) in Steinberg’s solution (58 mM NaCl, 0.67 mM KCl, 0.34 mM CaCl_2_, 0.85 mM MgSO_4_ and 4.6 mM Tris-HCl, pH 7.4) and centrifuged at 700*g* for 10 min at 20 °C. The centrifuged embryos were incubated in Steinberg’s solution for 1 h and transferred into CMFM. After incubating for 1 h, fertilization membranes were manually removed with fine forceps and translucent spherical blastomeres from the animal hemispheres were isolated with a glass rod in CMFM. The dissociated translucent blastomeres were subjected to fixation and immunostaining using the following procedures.

### Immunohistochemistry of dissociated blastomeres

The immunohistochemistry of dissociated blastomeres was performed as previously described (Nakamura et al, 2005) with slight modifications. The dissociated blastomeres were fixed in a fixative (80 mM piperazine-N,N’-bis (2-ethanesulfonic acid) [PIPES], 1 mM MgCl_2_, 5 mM ethylene glycol bis(2-aminoethyl ether)-N,N,N’,N’-tetraacetic acid [EGTA], 0.2% Triton X-100, 3.7% formaldehyde, 0.25% glutaraldehyde) overnight at room temperature, followed by washing with 0.1% TritonX-100 in Phosphate Buffered Saline (PBS: 268 µM KCl, 1.763 mM KH_2_PO_4_, 136.8 mM NaCl, 8.04 mM Na_2_HPO_4_, pH 7.4 (NaOH)) three times and post fixation in 100% methanol overnight at −20°C. After removing methanol through serial dilution using PBS containing 0.1% Tween-20 (PBST), the blastomeres were transferred to 0.2× SSC (75 mM NaCl, 7.5 mM Na-citrate) containing 2% H_2_O_2_ and 5% formamide for 2 h under light to bleach the pigments. After removing the H_2_O_2_ solution by washing with PBST twice, the blastomeres were incubated in blocking solution (PBS containing 2% bovine serum albumin and 0.1% TritonX-100) for 2 h at room temperature. The blastomeres were transferred to anti-NPC antibody (BioLegend, Clone: Mab414; 200× dilution) in blocking solution and incubated overnight at 4°C. After extensive washing with PBST (12 h, three times, at 4°C), the blastomeres were incubated with Alexa Fluor^®^ 488 conjugated anti-mouse IgG antibody (Thermo Fisher Scientific, 1752514; 200× dilution) in blocking solution overnight at 4°C. After extensive washing with PBST (12 h, three times, at 4°C), the blastomeres were stained with 1 µM TO-PRO-3 iodide (Thermo Fisher Scientific, T3605) for 30 min at room temperature. The blastomeres were washed with PBST three times and with 100% methanol three times. Finally, the blastomeres were transferred to a glass bottom dish (35 mm/Glass Base Dish, Iwaki) with a penetrating solution (benzyl alcohol:benzyl benzoate = 1:2). Using a wide-field microscope (Nikon, Eclipse Ti-E) equipped with 20× objective (Nikon, Plan Apo) and sCMOS camera (Andor, Zyla4.2) motorized by Micromanager or a confocal microscope (Olympus, FV1200) with 20× objectives (Olympus, UPlan SApo), images of nuclei in the blastomeres were acquired with z sections of 1 µm interval. The images acquired using the wide-field microscope processed were deconvoluted using the Microvolution ImageJ plugin (Microvolution, CA, USA).

### Cell-free reconstruction using *Xenopus* egg extracts

Crude cytostatic factor (CSF) metaphase-arrested extracts and cycling extracts from unfertilized *X. laevis* eggs were prepared as described in previous studies (Iwabuchi et al, 2000; Ohsumi et al, 2006). To reconstruct the interphase nuclei, CSF extracts were supplemented with 100 µg/ml cycloheximide, 1 µM tetramethyl-rhodamine-dUTP (TMR-dUTP: Roche, 11534378910; in the case to visualize DNA replication) and 0.6 mM CaCl_2_. Subsequently, demembranated sperm (150–200 /µl) was mixed with the extract and the samples were incubated for indicated time period at 22°C. Extracts showing the formation of nuclei with aberrant morphology were excluded from further experiments. To recapitulate cell cycle, the cycling extracts were supplemented with demembranated sperm and incubated at 22°C. The extracts containing sperm chromatin were fixed and stained by mixing with the same volume of 4% formaldehyde and 15% glycerol in extraction buffer (EB; 100 mM KCl, 5 mM MgCl_2_, 20 mM HEPES–KOH, pH 7.5) containing 5 µg/ml Hoechst 33342 (Invitrogen, H1399) and 10 µg/ml 3,3’-dihexyloxacarbocyanine iodide (DiOC_6_(3), Sigma, 318426). ∼4 µl of the mixture was put on a glass slide and covered with a coverslip. These observation slides were subjected to fluorescent imaging using the above-mentioned wide-field microscope equipped with a 40× objective lens (Nikon, Plan Fluor). For time-lapse imaging in the extracts, ∼10 µl of CSF extracts supplemented with CaCl_2,_ cycloheximide, sperm chromatin, 2 µM GFP-NLS recombinant proteins, and 1 µg/ml Hoechest 33343 was put on the glass slide between two lines of spacer tape (commercial 100-µm-thick double-sided tape) and covered by a coverslip with sealing the open parts with VALAP (mixture of Vaseline, lanolin, and paraffin). The sample on the imaging slide was incubated at 22°C on the above-mentioned wide-field microscope equipped with a 20× objective lens. To observe ER membranes based on a previous method (Wang et al., 2013), CSF extracts were prelabeled with 50 µg/ml of 1,10-dihexadecyl-3,3,3’,3’-tetramethylindocarbocyanine perchlorate (DiIC_16_(3); AAT Bioquest, 22044) for 45 min at 22°C. Reactions were prepared by adding 1:10 volume of pre-labeled extracts to a fresh CSF extracts mixed with CaCl_2,_ cycloheximide, sperm chromatin, and 1 µg/ml of Hoechest33343. Time-lapse imaging of the samples was carried out using the above-mentioned wide-field microscopy. For immunostaining of tubulin, extracts containing the reconstructed nuclei were fixed and stained as described previously (Hara and Merten, 2015) using an anti-α-tubulin antibody (DM1A clone; Thermo Fisher Scientific, MS581P0; 1,000× dilution), Alexa Fluor^**®**^ 633 conjugated anti-mouse IgG antibody (Thermo Fisher Scientific, A11003; 1,000× dilution). The samples were subjected to z-sectional imaging using the above-mentioned wide-field microscope equipped with 20× objective lens, and processed deconvolution using the Microvolution ImageJ plug-in.

### Isolation of yolk platelets

Based on a previous method (Wallace & Karasaki, 1963), yolk platelets were isolated from *X. laevis* unfertilized eggs. First, the eggs were dejellyed as described above and washed with 0.25 M sucrose in EB. The eggs were crushed using a Dounce tissue grinder (using a tight pestle; Wheaton, 357542) and centrifuged on a cushion of 1 M sucrose in EB at 600*g* and 4°C for 20 min. After washing the pellets with 0.25 M sucrose in EB once, the suspension of the pellet with 0.25 M sucrose in EB was centrifuged again on a cushion of 1 M sucrose in EB at 600*g* and 4°C for 20 min. The final pellets were diluted with EB at 1-2×10^6^/µl, which were used as the bulk fraction of the isolated yolk platelets. For further fractionation of the yolk platelets, the solution was centrifuged at ∼100*g* and 4°C for 20 min. The resulting supernatant was concentrated by further centrifugation at 600*g* to obtain a concentration of 1-2×10^6^/µl (small-sized fraction). The pellet was dissolved in EB to obtain a concentration of 1-2×10^6^/µl (large-sized fraction). The isolated yolk platelets were confirmed by staining with 0.5 mg/ml of Nile Red (Wako, 144-08811). When utilizing artificial beads instead of yolk platelets, magnetic beads (Veritastk, Dynabeads M-280: 3 µm diameter or Dynabeads T1: 1 µm diameter) at concentrations of 5×10^6^/µl (high) or 5×10^5^/µl (low) or polystyrene beads (Polysciences, 64090-15: 6 µm diameter) at concentrations of 1×10^5^/µl (high) or 1×10^4^/µl (low) were used. Before supplementation into the extracts, these artificial beads were washed with EB. The isolated yolk platelets or artificial beads were supplemented into cytoplasmic extracts at the same time as the supplementation of demembranated sperm chromatin. For the control, we supplemented with EB of the same volume.

### Quantification of nuclear size and cellular parameters

The cross-sectional area of the nuclei in the fixed blastomeres was measured using a ROI of Elliptical Sections in ImageJ software (National Institute of Health). For carrying out the measurement, we selected the z-plane that revealed the maximum nuclear cross-sectional area from the z series. Since this cross-sectional area can accurately calculate the nuclear surface area and volume of the dissociated blastomeres as a complete sphere as previously described (Edens and Levy, 2014), the nuclear and cell volumes were calculated as a complete sphere in this study. For measurement of the reconstructed nuclei on the fixed glass slide, the cross-sectional area was measured using an ellipsoid function in ImageJ software. The ROI of the Elliptical Sections was set on the rim of DiOC_6_(3)-positive nuclear membranes or along the inner side of the membrane accumulation around the DNA signal if the rim signal was not clear. For quantification of the nuclear expansion dynamics, we calculated the mean value of the cross-sectional area at each incubation time point and utilized the data set of the mean values in extract preparation. The values of at least 30 nuclei were measured for each experimental condition such as different time points and supplements. Each experimental condition was repeated using at least three individual extracts. To reduce the effects of variations in extract preparation, we have used only data showing more than 35 µm diameter in the reconstructed nuclei with *X. laevis* sperm chromatin without any supplements after 120 min of incubation for further analyses. Moreover, for normalization, the calculated mean values at a particular time point and condition (e.g., extract supplemented with yolk platelets for 80 min of incubation) were divided by the mean value obtained after 120 min of incubation in the control condition using exactly the same extract preparation (e.g., extract supplemented with buffer (EB) for 120 min of incubation). Mean values of the mean nuclear cross-sectional area as well as of the mean standard deviations from every individual experiment are illustrated in the graphs. For calculating the speed of nuclear expansion using the data set of mean nuclear cross-sectional area (Heijo et al, 2020), we calculated a slope of the regression line in the nuclear cross-sectional area against time from 60 min to 120 min of incubation in each extract preparation.

For quantification of MTs-occupied space, the edges of the MT-occupied space were detected by binarization of the images of visualized tubulins around the nucleus. We measured the sectional area enclosed by the detected edges using the Analyze Particles tool of ImageJ software. To quantify the area occupancy of yolk platelets or magnetic beads within the MT-occupied space, the yolk platelets or beads, which were non-specifically stained using DiOC_6_(3), were selected by binarization and the areas of the individual yolk platelets or beads were measured using the Analyze Particles tool of ImageJ software. After summing all the individual areas up, the total area of the yolk platelets or magnetic beads was divided by the area of MT-occupied space and termed as the area occupancy.

For estimation of cell cycle duration using dissociated blastomeres, we measured the whole cell cycle duration of a single cycle from completion of one cytokinesis to the next, as well as the mitotic duration of cytokinesis, from the initiation of furrow formation to the completion of the furrow (generation of two sister blastomeres). For estimation of interphase and mitosis durations in the cycling extract, we measured the duration of 1st interphase, from initiation of the extract incubation to NEBD, and of 1st mitosis, from NEBD to enclosing DiOC_6_(3)-positive membranes around the chromatin. Since there is a variation in cell cycle progression in each chromatin even in the same extract preparation, we classified the cell cycle only when more than half of the chromatin in the observation slide displayed membrane-enclosed (interphase) or membrane-less (mitosis) chromatin.

### Statistical analysis

We performed simple regression analysis by fitting regression lines in Excel software (Microsoft). To evaluate the strength of correlation, the coefficient of determination, R^2^ was calculated. Significant differences among samples were determined by non-parametric Wilcoxon tests using R software.

## ACKNOWLEDGEMENTS

We are grateful to Dr. Shuichi Ueno for suggestions to analyze the vegetal blastomeres and Dr. Wataru Kojima for supporting data analysis. *Xenopus tropicalis* was provided by Amphibian Research Center (Hiroshima University) through the National Bio-Resource Project of MEXT, Japan. This study was supported by the Home for Innovative Researchers and Academic Knowledge Users (HIRAKU) consortium, JSPS KAKENHI Grant Numbers JP16K14727, JP20H03253, and research grants from the Inamori Foundation and the Yamaguchi University Foundation to Y.H.

## AUTHOR CONTRIBUTIONS

S.S. and Y.H. performed most of the experiments and analyzed the data. Y.H with helpful supports from Y.I.

The authors declare no competing interest.

