## Supplemental figures for "Yolk platelets impede nuclear expansion in *Xenopus* embryos"

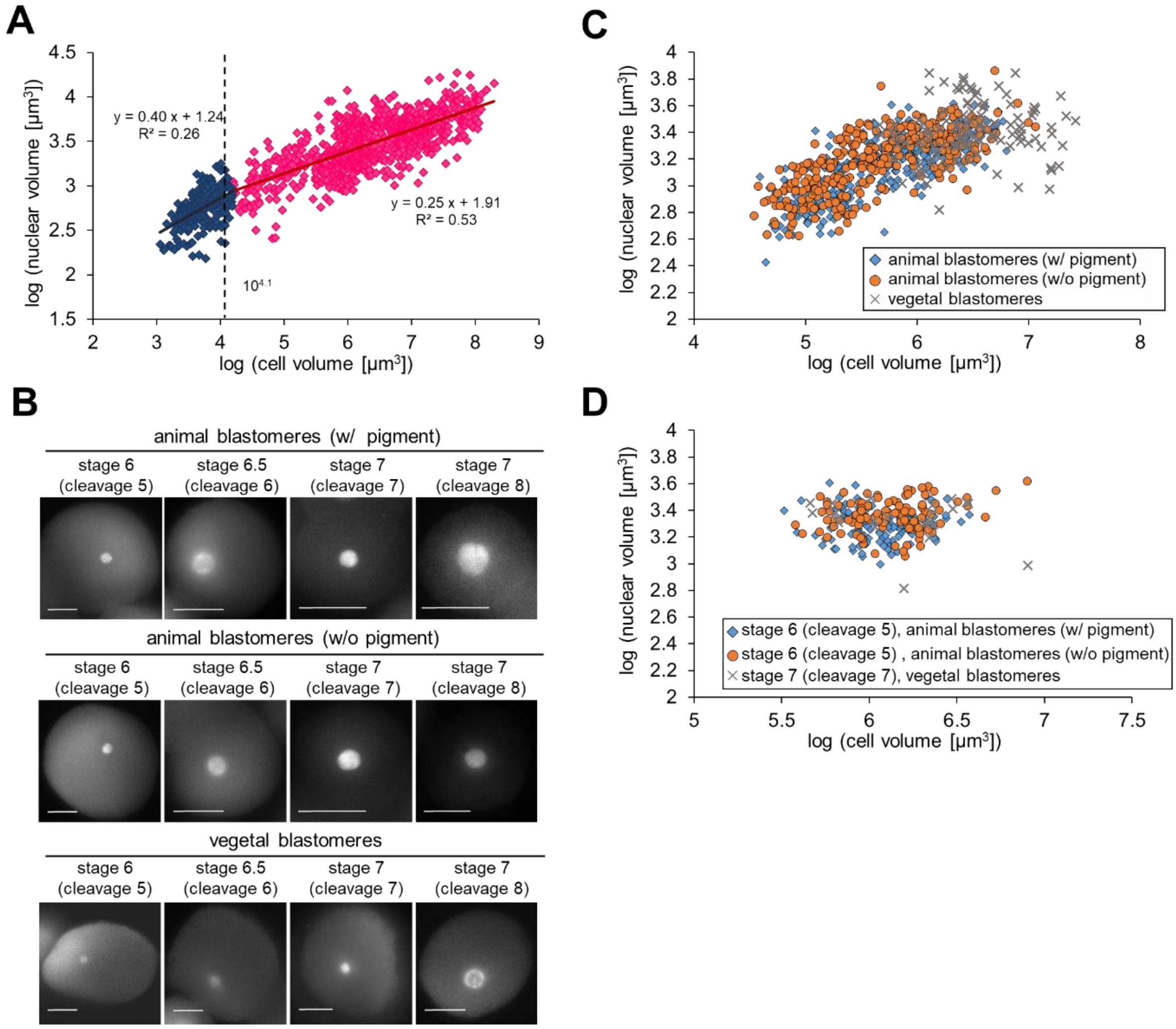


**Figure S1. Constant N/C ratio among animal and vegetal blastomeres from developing *X. laevis* and *X. tropicalis* embryos**

**(A)** Calculated nuclear volume was plotted against the calculated cell volume of blastomeres from different developmental stages of *X. laevis* embryos on a log-log plot. The data were identical to Fig. 1B and were fitted to liner regression with two segments (blue and pink lines). The segmented point was calculated using R software by evaluating the linearity and calculating the segmented point based on previous studies (Eberhard & Gutierrez, 1991; Hongo, 2007). After the calculation, the data within each group (more or less the segmented value) were fitted to liner regression using Excel (Microsoft) software. The dashed line indicates the border between the two segments. The cell volume of the border is ~13,000 μm$^3$. (**B**) DNA was stained using TO-PRO-3 in dissociated blastomeres from *X. tropicalis* developing embryos.

The blastomeres from embryos at various developmental stages were classified into three different groups; blastomeres with black pigments from the animal hemisphere [animal blastomeres (w/ pigments)], blastomeres without black pigments from the animal hemisphere [animal blastomeres (w/o pigments)], and blastomeres from vegetal hemispheres (vegetal blastomeres). Scale bar: 50 µm in stage 6 (cleavage 5)-7 (cleavage 7); 25 µm in stage 7 (cleavage 8). (**C**) Calculated nuclear volume was plotted against the calculated cell volume of blastomeres. The data of the nuclear and cell volumes were classified into the different blastomere groups (different colors). (**D**) Only data of the nuclear and cell volumes in animal blastomere from stage 6 (cleavage 5) and vegetal blastomeres from stage 7 (cleavage 7) *X. tropicalis* embryos were represented.

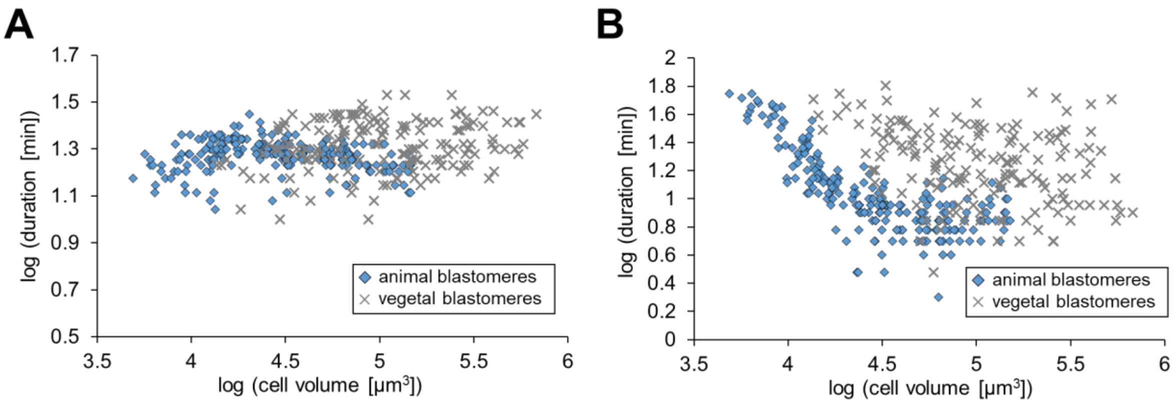


**Figure S2. Cell cycle duration in animal and vegetal blastomeres**

(**A**) Measured duration from initiation to completion of the of cleavage furrow formation furrow was plotted against the calculated mean cell volume during the cell cycle. (**B**) Calculated duration, in which the cytokinesis duration was subtracted from duration for cell cycle from the completion of one cytokinesis to the next, was plotted against the mean cell volume. The data were acquired from each dissociated animal (blue) or vegetal (grey) blastomeres.

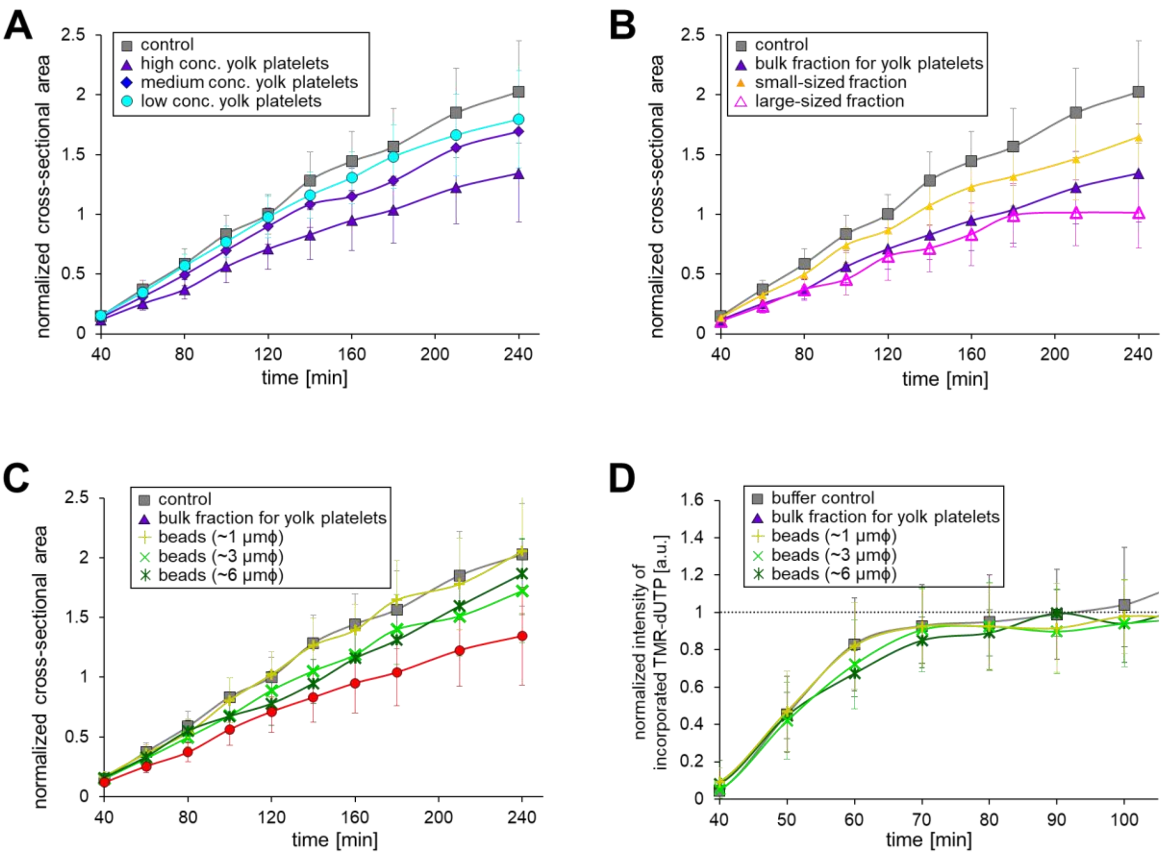


**Figure S3. Supplementation with yolk platelets or artificial beads in cell-free extracts decreases the speeds of interphase progression**

(**A**) Dynamics of the mean normalized cross-sectional area in the nuclei reconstructed with different concentrations of yolk platelets. (**B**) Dynamics of the mean normalized cross-sectional area in the reconstructed nuclei by supplementation with different-sized yolk platelets. (**C**) Dynamics of the mean normalized cross-sectional area in the reconstructed nuclei upon supplementation with each artificial beads. (**D**) Dynamics of the intensities of the incorporated TMR-dUTP in whole nuclei in the presence of artificial beads with different sizes. The intensity was calculated by multiplying the measured TMR-dUTP intensity (/$\mu m^2$) with the measured nuclear cross-sectional area. Each calculated value was divided by the mean value of the individual extract preparation after 120 min of incubation in the absence of any supplements. The symbols represent the mean normalized nuclear cross-sectional area or TMR-dUTP intensity from fixed sample. Averages of values are connected by a line in each dataset. Error bar: SD.
